# S100A1’s single cysteine is an indispensable redox-switch for the protection against diastolic calcium leakage in cardiomyocytes

**DOI:** 10.1101/2023.09.01.555853

**Authors:** Andreas Seitz, Martin Busch, Jasmin Kroemer, Andrea Schneider, Stephanie Simon, Andreas Jungmann, Hugo A. Katus, Julia Ritterhoff, Patrick Most

## Abstract

The EF-hand calcium (Ca^2+^) sensor protein S100A1 combines inotropic with antiarrhythmic potency in cardiomyocytes (CM). Oxidative posttranslational modification (ox-PTM) of S100A1’s conserved, single cysteine residue (C85) via reactive nitrogen species (i.e. S-nitrosylation or glutathionylation) was proposed to modulate conformational flexibility of intrinsically disordered sequence fragments and to increase the molecule’s affinity towards Ca^2+^. In light of the unknown biological functional consequence, we aimed to determine the impact of the C85 moiety of S100A1 as a potential redox-switch.

We first uncovered that S100A1 is endogenously glutathionylated in the adult heart in vivo. To prevent glutathionylation of S100A1, we generated S100A1 variants that were unresponsive to ox-PTMs. Overexpression of wildtype (WT) and C85-deficient S100A1 protein variants in isolated CM demonstrated equal inotropic potency, as shown by equally augmented Ca^2+^ transient amplitudes under basal conditions and β-adrenergic receptor (βAR) stimulation. However, in contrast ox-PTM defective S100A1 variants failed to protect against arrhythmogenic diastolic sarcoplasmic reticulum (SR) Ca^2+^ leak and ryanodine receptor (RyR2) hypernitrosylation during β-AR stimulation. Despite diastolic performance failure, C85-deficient S100A1 protein variants exerted similar Ca^2+^-dependent interaction with the RyR2 than WT-S100A1.

Dissecting S100A1’s molecular structure-function relationship, our data indicate for the first time that the conserved C85 residue potentially acts as a redox-switch that is indispensable for S100A1’s antiarrhythmic but not its inotropic potency in CM. We therefore propose a model where C85’s ox-PTM determines S100A1’s ability to beneficially control diastolic but not systolic RyR2 activity.

## 1. Introduction

S100A1 is an EF-hand calcium (Ca^2+^) sensor protein highly expressed in the heart [1]. Molecular studies utilizing both gene-deletion as well as gene-addition strategies characterized S100A1 as a positive regulator of cardiomyocyte (CM) performance [2-7]. Lack of S100A1 expression in cardiomyocytes (CM) disabled the mammalian heart to cope both with acute and chronic hemodynamic stress, and results in accelerated transition to contractile failure [4]. By targeting the activity of numerous downstream key effector proteins of the sarcoplasmic reticulum (SR) (RyR2 and SERCA2a), the contractile apparatus (cardiac titin) and mitochondria (Complex V), S100A1 plays a critical role as an upstream regulator of the Ca^2+^-controlled contraction-relaxation cycle, energy homeostasis, stress resilience and rhythm stability of CM [2, 6, 8, 9]. A direct interaction between S100A1 and its various target proteins is a prerequisite for beneficial activity modulation, which is facilitated by the Ca^2+^ activated “on-state” of the EF-hand Ca^2+^ sensor [3, 6].

Structural guided molecular imaging studies raised important questions about the impact of posttranslational modifications (PTMs) on the structure-function relationship of the S100A1 homodimer [10-12]. In particular, S100A1’s conserved single cysteine residue 85 (C85), located in the C-terminal α-helix of each monomer, attracted attention due to its high reactivity towards oxidative PTMs (ox-PTMs). Here, S-nitrosylation (SNO) or S-glutathionylation (SSG) of C85 emerged as most prevalent PTMs. Subsequently entailing conformational changes, both Ca^2+^ affinity of the two EF-hand domains are enhanced by several orders of magnitude and the structure of epitopes implicated in target binding are altered [12-15].

No study, however, has yet tested the relevance of C85 ox-PTM for S100A1’s biological function in cardiomyocytes. Given the therapeutic impact of an S100A1 gene-based therapy, we thus aimed to unveil the role of S100A1’s C85 residue as a potential regulatory redox-element in CM. To this end, we tested the hypothesis that C85 allows S100A1 to function as a redox-sensitive Ca^2+^ sensor in CM, thereby transitioning to a Ca^2+^ activated “on-state” already at low diastolic Ca^2+^ concentrations. Here, we demonstrate that S100A1 is endogenously modified by ox-PTMs and specifically, glutathionylated in the adult heart. We then generated S100A1 protein variants that are unresponsive to ox-PTMs at this site. We demonstrate that C85 is dispensable for enhancing systolic Ca^2+^ fluxes, but indispensable for the protection against diastolic SR Ca^2+^ leakage and RyR2 hypernitrosylation in CM upon catecholamine stress.

Such data advance our understanding of S100A1’s molecular function and, most likely, differentiate for the first time between diastolic redox-sensitive and systolic redox-insensitive molecular mechanisms that are expected to be of equal relevance for S100A1’s therapeutic efficacy in human heart failure.

## 2. Materials and Methods

### Experimental Animals

All animal procedures and experiments were performed in accordance with the ethical standards laid down in the 1964 Declaration of Helsinki and its later amendments institutional, guidelines of the University of Heidelberg and received approval from local authorities.

### Materials

Following substances and kits were used: DyLight 680 Maleimide (Thermo Fisher Scientific, 46618), Fura2-AM (life technologies, F-1225), Laminin (Sigma Aldrich, L2020), Medium M199 (Sigma Aldrich, M7528), Pierce Classic Magnetic IP/Co-IP Kit (Thermo Scientific, 88804), Biotin-maleimide (B1267, Sigma Aldrich), N-Ethylmaleimide (E3876, Sigma Aldrich), ROS-Glo H2O2 Assay (Promega, G8820). All other chemicals were obtained from Sigma Aldrich, Germany, if not indicated specifically.

### Generation of control and S100A1 variant adenoviruses

First-generation early gene 1/3-deleted adenoviruses were obtained by the use of the pAdTrack CMV/pAdEasy-1 system as previously described [2, 16]. Adenoviruses with C85A and C85S were generated as the wild-type S100A1 adenovirus. Expression of S100A1, C85A or C85S and green fluorescent protein (GFP) reporter gene was driven by two independent cytomegalovirus (CMV) promoters. The same adenovirus devoid of S100A1 cDNA served as control (AdGFP). A detailed description is available in the Supplemental Material and Methods.

### Modified Biotin switch assay

A modified Biotin switch assay (mBSA) was performed with modifications from [17-19] as described in the Supplemental Material and Methods.

### Neonatal and adult rat cardiomyocyte isolation

Neonatal rat heart cells were isolated by gradual trypsin digestion from 1-3 days old neonatal Wistar rat pubs as described previously [4]. Isolated cells were pre-plated to enrich cardiomyocyte fraction. Cardiomyocytes were plated with density of 1 Mio cells/ml and cultured for 3 days before viral transduction in medium M199 supplemented with 10%FCS (day 1-2)/0.5% FCS (from day 3) and 10,000 U/ml penicillin/10 mg/ml streptomycin, 1% l-glutamine, 1mM CaCal_2_.

Ca^2+^ tolerant adult rat cardiomyocytes were isolated from LV’s male Wistar rats, by a standard enzymatic digestion procedure and cultured as described previously [2]. Cardiomyocytes were plated with a density of 50.000 cells/ml on laminin-coated (1:100) fluorodishes (World Precision Instruments, USA) or culture plates. Adenoviral transduction of isolated ventricular cardiomyocytes was carried out 2 h after plating in HEPES-modified medium 199 (M199 supplemented with 5 mM taurine, 5 mM carnitine, 5 mM creatine, 5 mM N-mercaptoproprionyl glycine, 0.1 µM insulin, 10,000 U/ml penicillin and 10 mg/ml streptomycin, pH 7.25).

### Intracellular Ca^2+^ transients

Neonatal or adult rat cardiomyocytes were incubated with of Fura 2-AM (1 µM for 20 minutes at 37°C followed by 5 minutes incubation to allow for complete desterification of the dye) and measurements were carried out using an inverse Olympus microscope (IX70) with a UV filter connected to a monochromator (Polychrome II, T.I.L.L. Photonics GmbH, Germany) as previously described [2, 20, 21]. Cells were electrically stimulated with a biphasic pulse to contract at room temperature and excited at 340/380 nm. Epifluorescence emission was detected at 510 nm, digitized, and analyzed off-line with T.I.L.L.VISION software (v. 3.3). Isoproterenol (1µM) was added after baseline recording when indicated and measurements repeated after 5 minutes incubation.

Data from 5 consecutive steady-state transients were averaged for analysis of transient amplitude using a personal automatic detection of transients.

### Sarcoplasmic reticulum (SR) Ca^2+^ leak assay: Rapid pacing

For the rapid pacing, adult rat cardiomyocytes were treated as described above. Cardiomyocytes were stimulated for steady states Ca^2+^ transients at 3Hz for 20 seconds alternated by periods of rest of equal duration to allow for the occurrence of spontaneous diastolic Ca^2+^ waves. Isoproterenol stimulation (1µM) was added to an individual group of CM and experiments were performed after 5 minutes of incubation. Ca^2+^ waves were quantified manually by analyzing the number of cardiomyocytes with diastolic Ca^2+^ waves as well as the number of Ca^2+^ waves per cell. Ca^2+^ transient amplitude was analyzed as described above.

### Proximity Ligation Assay (PLA), Western Blotting and Immunoprecipitation

Proximity Ligation Assay (PLA), Western Blotting and Immunoprecipitation were performed as described previously and detailed description is available in the Supplemental Material and Methods [21].

### Statistics

The numbers of independent experiments are specified in the relevant figure legends. Data are expressed as mean + standard error of the mean (SEM). Statistical analysis was performed with Prism 9.0 software (GraphPad). Statistical comparisons between 2 groups were performed by unpaired, two-tailed t-test. Statistical comparisons between 3 or more groups were performed by one-way ANOVA followed by the appropriate posthoc analysis to determine statistical significance. Details are given in the respective figure legends. The value of p < 0.05 was considered statistically significant.

## 3. Results

### 3.1 S100A1 is endogenously s-glutathionylated in the adult heart

Biochemical studies suggested that S100A1 can be modified by ox-PTM, including s-nitrosylation (SNO) and s-glutathionylation (SSG) [10-12]. However, these modifications have never been demonstrated in cardiac tissue. To determine if S100A1 is endogenously modified in the heart in vivo, we used a modified biotin switch assay (mBSA) followed by immunoprecipitation of S100A1 (Suppl. Fig. 1) to detect oxidized cysteine modifications, including SNO and SSG [17-19]. S100A1 was successfully immunoprecipitated after mBSA, and endogenously modified by ox-PTM (Fig. 1A). Surprisingly, we could not detect endogenous SNO of S100A1, as it had previously been proposed in a non-cardiac cell type [12], but S100A1-SSG appeared as the prevalent modification (Fig. 1A).

**Fig. 1:**
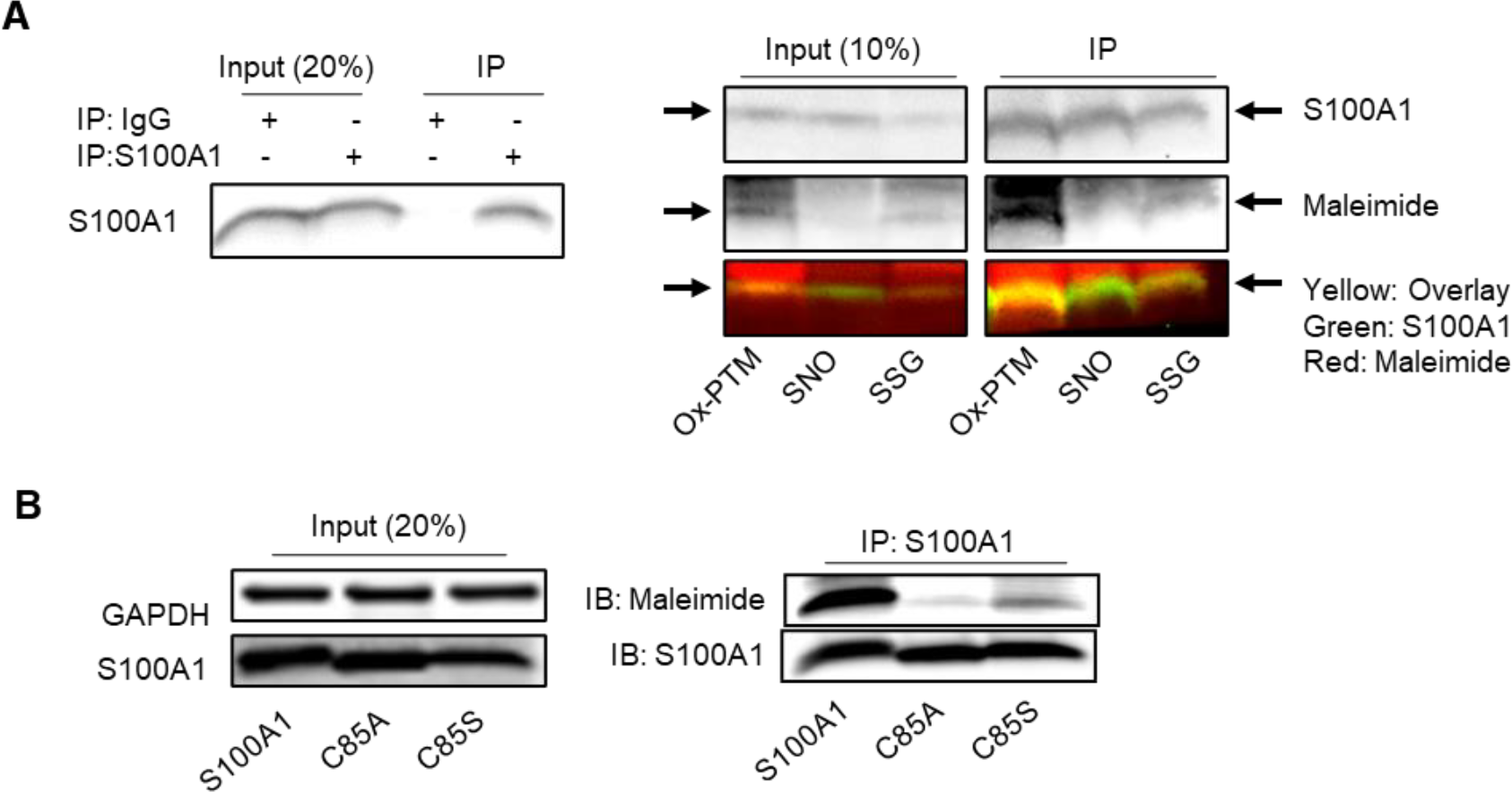
S100A1 is endogenously glutathionylated in vivo. A: left: Immunoprecipitation of S100A1 from mouse heart tissue demonstrating specificity of S100A1-IP. Right: Input and S100A1 IP after mBSA to detect ox-PTM (reduced with DTT), SNO (reduced with ascorbate) and SSG (reduced by Grx enzymatic reaction). The yellow overlay indicates that S100A1 is modified by ox-PTM and SSG. B: Representative immunoblot of maleimide incorporation after immunoprecipitation of WT- and mutated-S100A1 protein from HEK293T cells. Cell lysates were stained with maleimide to label reactive cysteine residues after control, S100A1, C85A or C85S overexpression. Input (20%) for S100A1 and GAPDH as loading control is depicted on the left side ensuring matching S100A1 overexpression.

To test the functional relevance of this modification, we then generated adenoviruses (Ad) that express human S100A1 protein variants resistant to ox-PTM (C85A and C85S). Functional lack of reactive cysteines was demonstrated by overexpression WT and C85-deficient S100A1 variants and labeling with maleimide, which specifically interacts with free thiols. After immunoprecipitation of S100A1, only WT-S100A1 but not C85-deficient S100A1 protein variants were labeled with maleimide (Fig. 1B). Of note, a minor fluorescent signal was still visible both in C85A- and C85S-S100A1 overexpressing CM indicative of endogenous WT-S100A1 protein.

Together, this demonstrates that glutathionylation is the predominant modification of S100A1 in the heart, and that C85-deficient S100A1 lack functional thiols.

### 3.2 C85A- and C85S-S100A1 increase systolic Ca^2+^ transients in CM

We next assessed the functional impact of C85-deficient S100A1 protein variants on systolic intracellular Ca^2+^ amplitudes in isolated adult rat cardiomyocytes (ARCM).

Adenoviral delivery of WT-as well as C85A- and C85S-S100A1 resulted in an approximately 5-fold increase of each S100A1 protein variant after 24 hours (Fig. 2A, Supplemental Fig. S2A, B). Of note, overexpression levels matched those achieved in previous studies that disclosed the inotropic actions of S100A1 in rodent CM [21-23]. All overexpressed S100A1 protein variants yielded a significant but indistinguishable increase in Ca^2+^ transient amplitudes under basal conditions (Fig. 2B, C). Alike WT-S100A1, the gain-of-function by C85-deficient S100A1 variants was preserved under βAR stimulation using isoproterenol (ISO) (Fig. 2B, C).

**Fig. 2:**
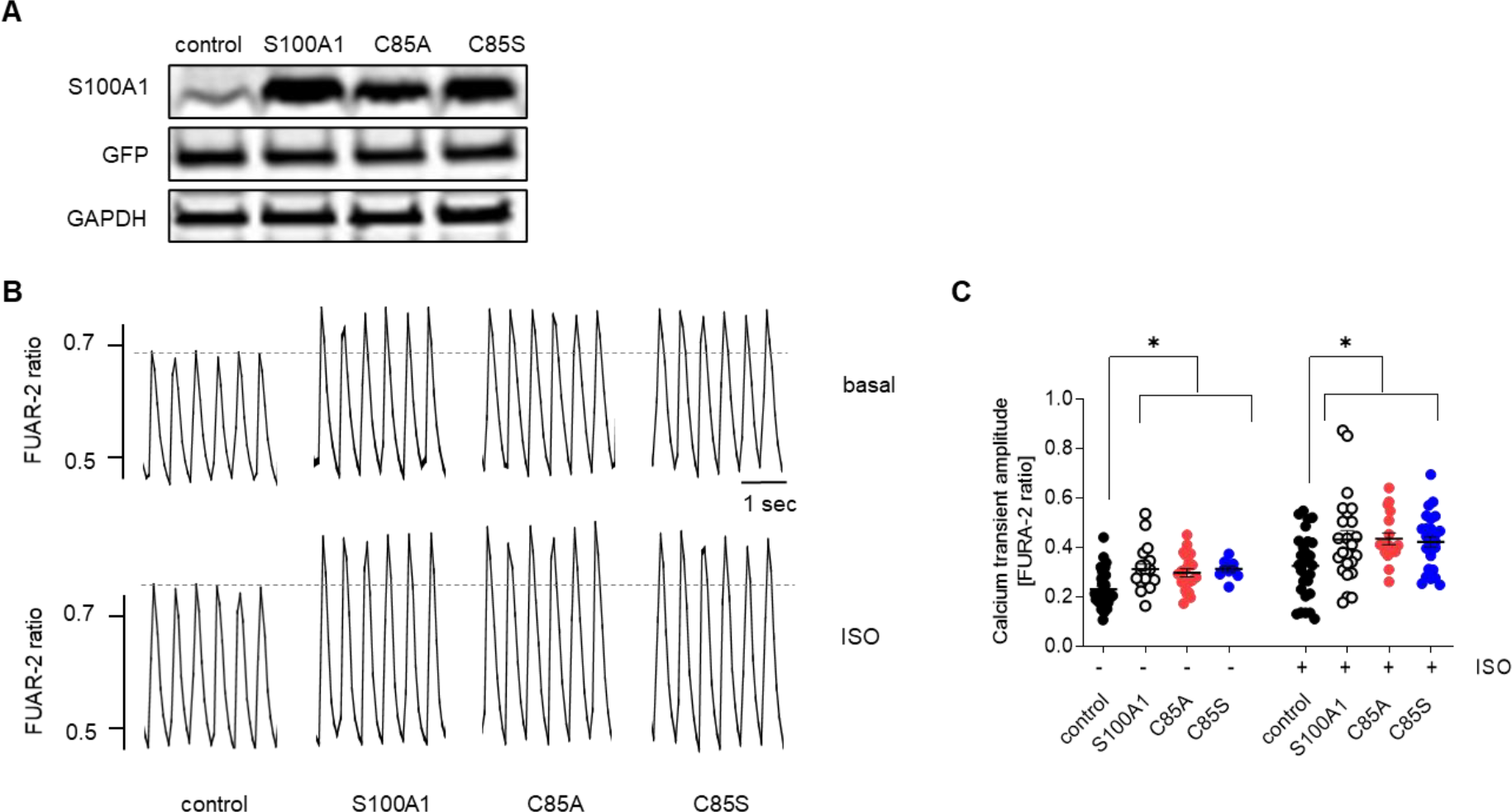
WT and C85-deficient S100A1 overexpression increases systolic Ca^2+^ transient amplitudes in CMs. **A:** Representative immunoblots from control, S100A1, C85A and C85S treated adult rat cardiomyocytes (ARCM) 24h after adenoviral transduction demonstrating equal S100A1 overexpression and viral load. **B:** Representative tracings of electrically-stimulated (2Hz) steady-state Ca^2+^ transients from control, S100A1, C85A and C85S overexpressing ARCM. **C:** Quantification of B. Overexpression of WT and C85-deficient S100A1 results in increased calcium transients. n = 20-40; cells in each group were derived from 3 different isolations; 1-way ANOVA with Dunnett’s multiple comparison post hoc test, *p <0.05. Data are given as mean±SEM.

Thus, S100A1’s C85 residue seems dispensable for increasing systolic Ca^2+^ transient amplitudes in CM.

### 3.3 C85-deficient S100A1 variants fail to protect βAR-triggered diastolic arrhythmogenic sarcoplasmic reticulum Ca^2+^ waves

Given the previously reported protective effect of overexpressed WT-S100A1 against diastolic RyR2 dysfunction and arrhythmogenic SR Ca^2+^ leakage [21], we subsequently determined efficacy of C85-deficient S100A1 protein variants on diastolic CM performance. To this end, we employed a previously published steady-state rapid pacing (3Hz) protocol to enhance SR Ca^2+^ load, which enables formation of diastolic Ca^2+^ waves due to abrupt cessation of the stimulation (Fig. 3A).

**Fig. 3:**
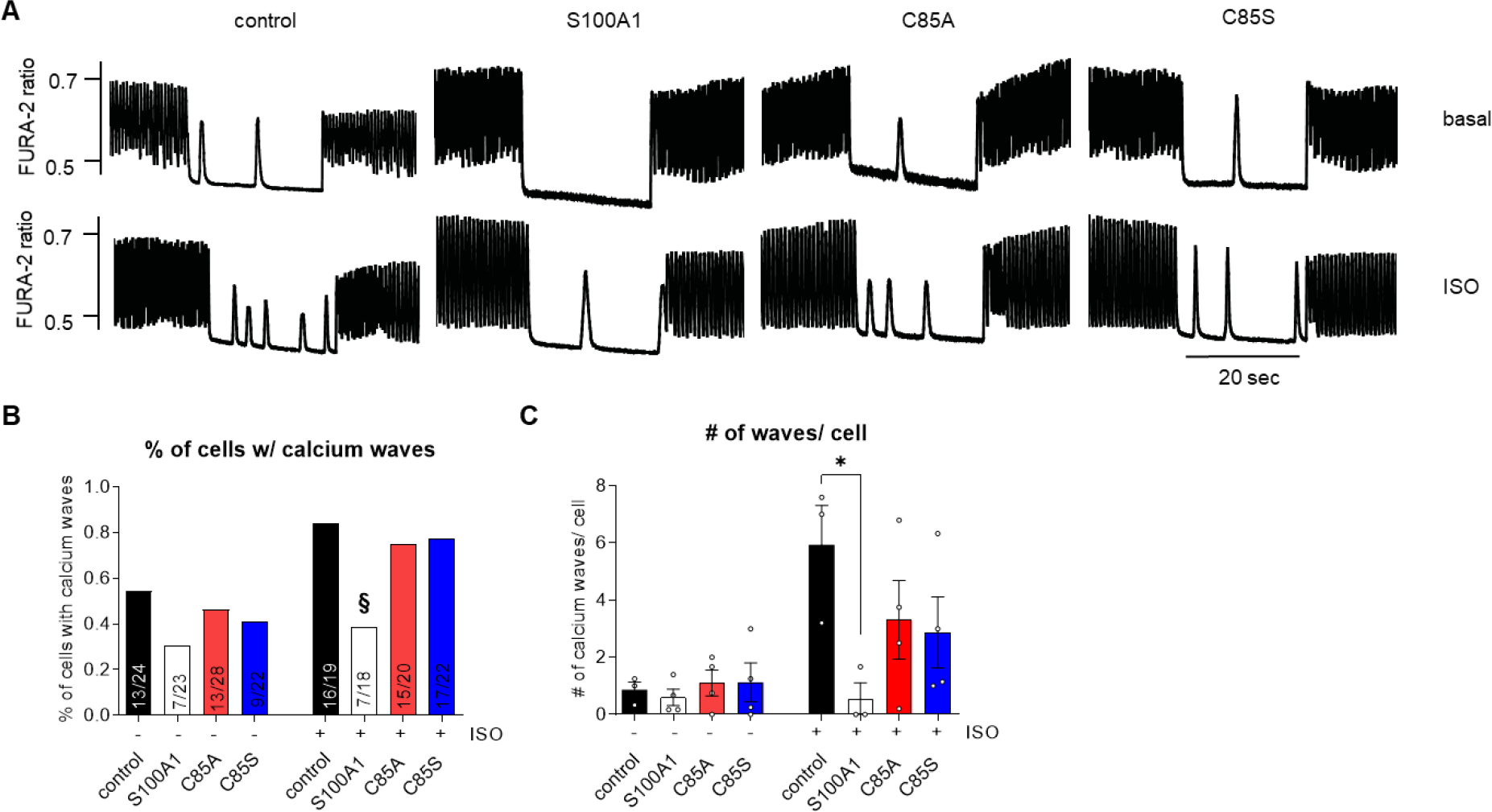
WT- but not C85-deficient S100A1 protects against rapid pacing-induced calcium waves in CMs. **A:** Representative tracings of electrically-stimulated Ca^2+^ transients from basal and ISO-stimulated (1 µM) control, S100A1, C85A or C85S treated ARCM. Cells were electrically stimulated (3Hz) for 20 sec followed by a 20 sec resting period and restimulation (3Hz) for 20 sec. **B:** The percentage of cells with spontaneous calcium waves were quantified ISO stimulation results in an increase in calcium waves in control, C85A and C85S treated cardiomyocytes but not after WT S100A1 overexpression. Number of cells analyzed per group is shown in each bar. Cells in each group were derived from 4 different isolations; Chi Square test/Fisher’s exact test. §p<0.05. **C:** Number of calcium waves per cell were quantified ISO stimulation results in an increase in calcium waves in control, C85A and C85S treated cardiomyocytes but not after WT S100A1 overexpression. n = 3-4. 1-way ANOVA with Dunnett’s multiple comparison post hoc test. * p<0.05. Data are given as mean±SEM.

Under baseline conditions, WT-S100A1 tended to decrease the incidence of the SR Ca^2+^ leak, although this trend did not reach statistical significance (Fig. 3A-C). As desired, βAR stimulation by ISO exaggerated the incidence of spontaneous SR Ca^2+^ release that was measured both by the number of CM with diastolic Ca^2+^ release and number of Ca^2+^ waves per cell (Fig. 3A-C). Overexpression of WT-S100A1 completely prevented the βAR-mediated increase in diastolic Ca^2+^ waves in CM compared to controls, which concurs with previous results [21]. Although C85A- and C85S-S100A1 protein variants also increased the systolic Ca^2+^ transient amplitudes during the 3Hz steady-state pacing period to the same extent than WT-S100A1, both C85-deficient S100A1 protein variants failed to protect CM against spontaneous and βAR-triggered diastolic Ca^2+^ waves (Fig. 3A-C).

Hence, C85 appears indispensable for S100A1’s control of the RyR2 of diastolic leakage.

### 3.4 C85A- and C85S-S100A1 interact with the RyR2 at diastolic Ca^2+^ levels

Failure of C85-deficient S100A1 variants to prevent βAR-triggered RyR2 dysfunction prompted the question whether C85 substitution might disable S100A1 to interact with the RyR2 and/or unfavorably alter posttranslational modification of the SR Ca^2+^ release channel.

As previous reports linked PKA and CamKII-mediated RyR2 hyperphosphorylation to diastolic gating dysfunction [24, 25], we next determined RyR2 phosphorylation at the corresponding serine-2808 and -2814 sites (Fig. 4A). However, none of the overexpressed S100A1 variants mitigated the βAR-induced increase in RyR2 phosphorylation (Fig. 4A, B). Additionally, phospholamban (PLB) was included to capture potential differential activity on βAR downstream signaling in other SR compartments. Again, no differences between the overexpressed S100A1 protein variants on global signaling were observed (Fig. 4A, C).

**Fig. 4:**
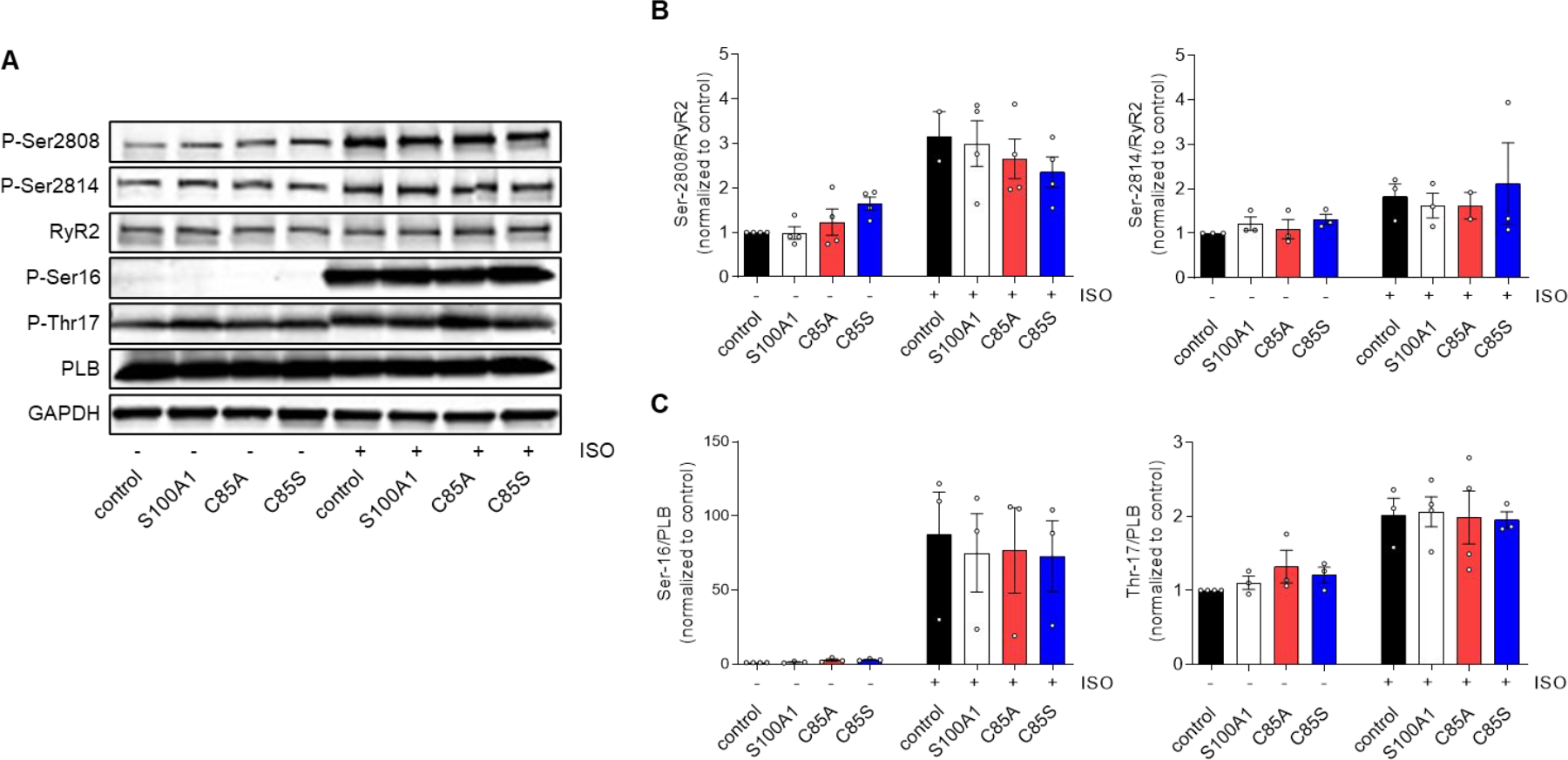
WT- and C85-deficient S100A1 overexpression has no impact on RyR2 and PLB phosphorylation in CMs. **A:** Representative immunoblots for total PLB, PLB Ser-16 and Thr-17, total RyR2, RyR Ser-2808 and Ser-2814 in control, S100A1, C85A and C85S treated ARCM. Quantification shows no differences in RyR2 (**B**) and PLB (**C**) phosphorylation levels between control, S100A1, C85A and C85S treated CMs, whereas ISO stimulation increases PKA-/CamKII phosphorylation of RyR2 and PLB. n=4; 1-way ANOVA with Dunnett’s multiple comparison post hoc test, * p<0.05. Data are given as mean±SEM.

Furthermore, ox-PTM of C85 might be required for S100A1’s Ca^2+^-dependent interaction with target proteins [6]. We thus assessed the capability of overexpressed S100A1 variants to interact with the RyR2 in neonatal rat cardiomyocytes (NRCM), which exhibited the same inotropic response after overexpression of the S100A1 protein variants (Suppl. Fig. 3A-D). Co-immunoprecipitation analysis demonstrated that C85A- and C85S-S100A1 increased the S100A1/RyR2 ratio as WT-S100A1 under high Ca^2+^ conditions (Fig. 5A). We subsequently used the in-cell proximity ligation assay (PLA) to study S100A1/RyR2 binding, as this method enables the detection of protein-protein interactions within their intact subcellular domains at near diastolic Ca^2+^ concentrations in quiescent CM (Fig. 5B). Interestingly, all overexpressed S100A1 protein variants resulted in a significant 2-fold increase of the S100A1/RyR2 ratio at baseline and after βAR stimulation (Fig. 5B, C).

**Fig. 5:**
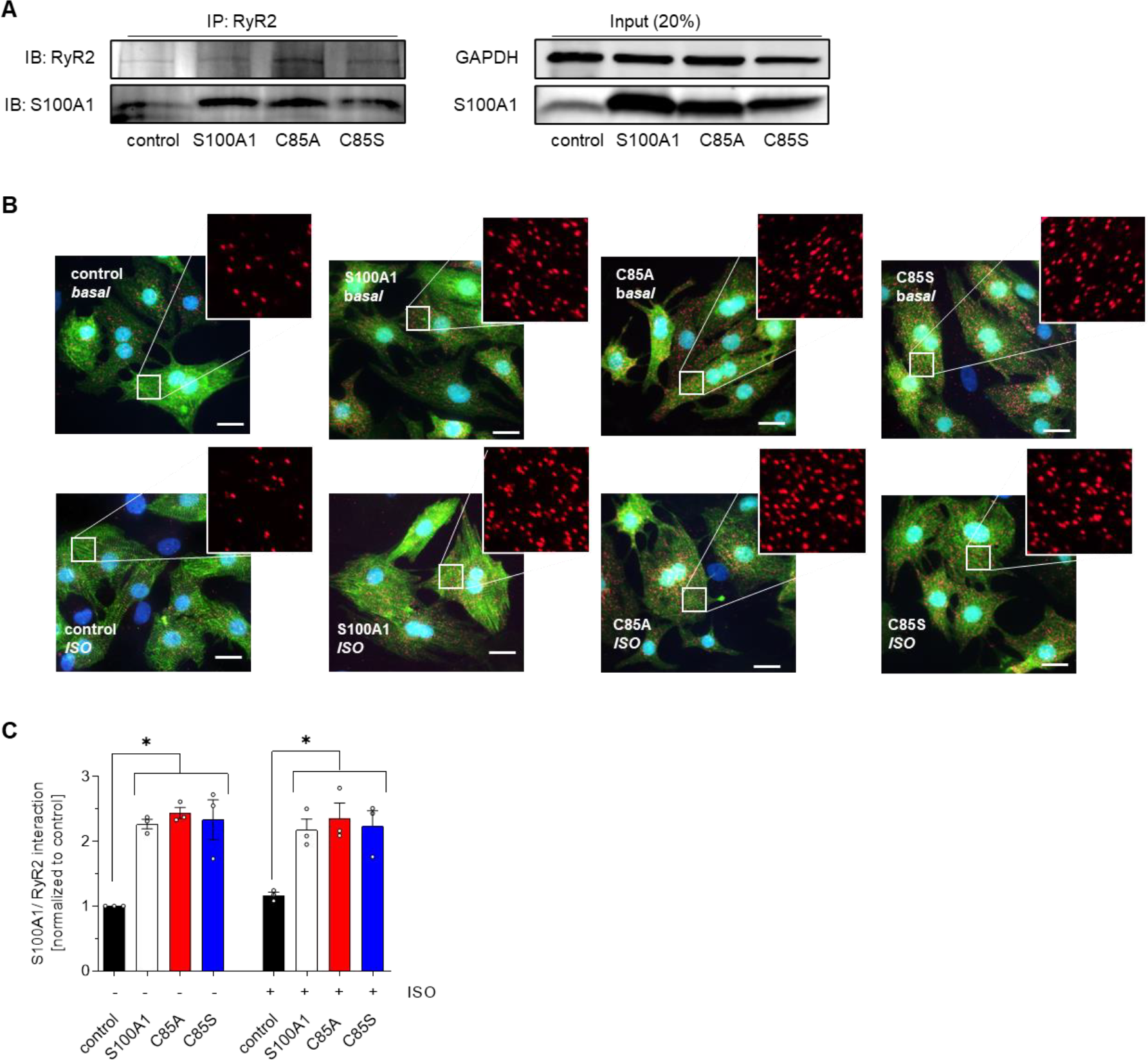
WT- and C85-deficient S100A1 interact with the RyR2. **A:** Representative immunoblot for RyR2 and co-precipitating S100A1 protein after RyR2 immunoprecipitation from control, S100A1, C85A or C85S treated neonatal rat cardiomyocytes (NRCM). Left: Input (20%) for S100A1 and GAPDH as loading control. **B:** Representative proximity ligation assay (PLA) images from control, S100A1, C85A and C85S treated NRCM showing S100A1/RyR2 interaction (red dots). Inlet magnification is 2-fold. Scale bar represents 20 µm. **C:** Statistical analysis shows a 2-fold increase of the S100A1/RyR2 binding ratio. n=4; 1-way ANOVA with Dunnett’s multiple comparison post hoc test, * p<0.05. Data are given as mean±SEM.

Together, this demonstrates that C85-deficient S100A1 did not change the S100A1/RyR2 stoichiometry in quiescent CM at systolic and diastolic Ca^2+^ levels, despite persistent altered RyR2 phosphorylation.

### 3.5 C85-deficient S100A1 variants fail to abrogate βAR-induced RyR2 hypernitrosylation

Besides phosphorylation, ßAR stress has been reported to render the SR Ca^2+^ channels leaky via aberrant hypernitrosylation of distinct reactive cysteines [26, 27]. To determine if RyR2-SNO is affected by S100A1, we again employed the in-cell PLA assay. βAR-stimulation resulted in a significant increase of RyR2-SNO in control cells (Fig. 6A, B), as previously demonstrated [26, 28], and WT-S100A1 overexpression suppressed this βAR-mediated RyR2 hypernitrosylation. In contrast, both C85A- and C85S-S100A1 failed to abrogate hypernitrosylation of RyR2 under βAR stimulation (Fig. 6A, B). Hence, this is the first evidence that S100A1 can impact the function of a crucial intracellular SR target by modulating the NO-dependent redox-state via its conserved C85 residue. NOS1 (nNOS)-mediated RyR2-SNO has been shown to modulate cardiac contractility and ventricular arrhythmias [28, 29], so we next investigated a potential role for S100A1 in compartmentalized SR NO-signaling. Here, inhibition of NOS1 with SMLT was sufficient to abrogate ISO-induced RyR2-SNO, but did not further reduce RyR2-SNO after WT-S100A1 overexpression (Fig. 6C), supporting the role of NOS1-mediated RyR2 hypernitrosylation. S100A1 has been demonstrated to interact with NOS3 (eNOS) in endothelial cells thereby modifying vascular NO-signaling [30]. Since NOS1 rather than NOS3 is targeted to cardiac SR and RyR2 [28], we sought to determine whether WT-S100A1 might interact with NOS1 or modulate its expression. However, we could not provide evidence for WT-S100A1 to interact with NOS1 in CM assessed by PLA or co-immunoprecipitation (data not shown) and thus could not establish a direct link between S100A1 and NOS1 activation.

**Fig. 6:**
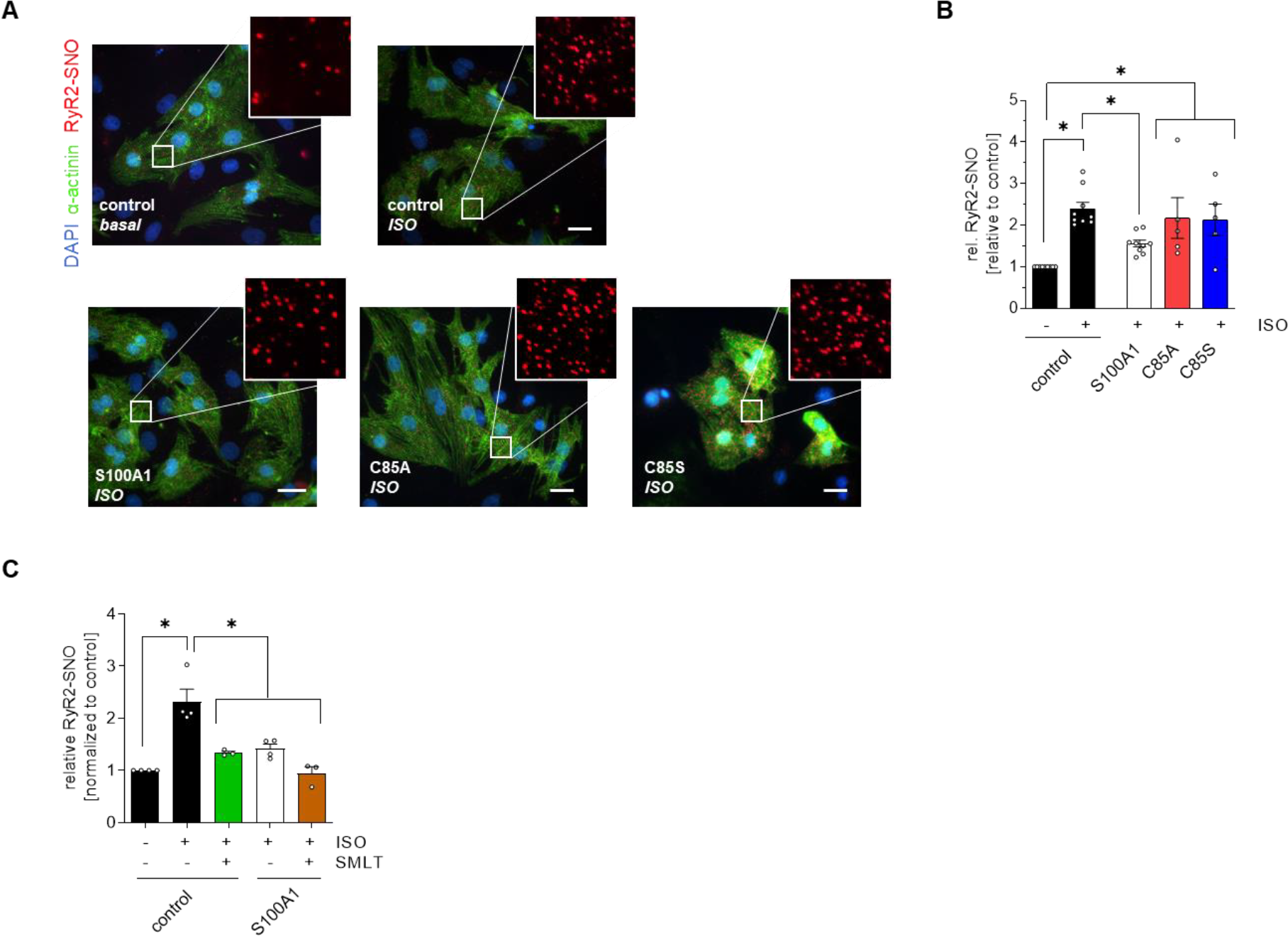
WT-S100A1 prevents RyR2 hypernitrosylation during β-adrenergic receptor stimulation. **A:** Representative PLA images from control, S100A1, C85A and C85S treated NRCM showing pan-RyR2 nitrosylation (red dots). Inlet magnification is 2-fold. Scale bar represents 20µm. B: Quantification of A. RyR2 is hypernitrosylated after ISO stimulation, which is prevented after WT, but not after C85-deficient S100A1 protein overexpression. n= 5-9, 1-way ANOVA with Tukey’s multiple comparison post hoc test, *p<0.05. C: Quantifaction of RyR2-SNO after SMLT stimulation in indicated groups. SMLT reduced RyR2-SNO during ISO stimulation in control, but not S100A1-treated CM. n = 3-4. 1-way ANOVA with Tukey post hoc test. Data are given as mean±SEM.

Alternatively to a direct, NOS1-dependent nitrosylation modification, a second pathway that increases oxidative stress in the vicinity of the target has been proposed to regulate nitrosylation of SR proteins [28]. To test this alternative hypothesis and assess if S100A1 affects whole-cell redox homeostasis, we measured cellular ROS generation in response to ISO stimulation in NRCM. Neither WT-, C85A-nor C85S-S100A1 overexpression had an impact on global ROS levels (Supplemental Fig. 4A), indicating that βAR stimulation did not evoke cellular ROS but specifically modulated RyR2-SNO. Thus, overexpressed WT-S100A1 protein might not generally act as a buffer for reactive oxygen species via C85 but exert compartmentalized actions confined to S100A1 target proteins.

Together, this let us to conclude that during diastole, under low Ca^2+^ conditions, S100A1 interaction with the RyR2 prevents RyR2 hypernitrosylation and subsequent SR Ca^2+^ leakage during βAR stimulation, which the later one being dependent on C85 of S100A1 (Fig. 7).

**Fig. 7:**
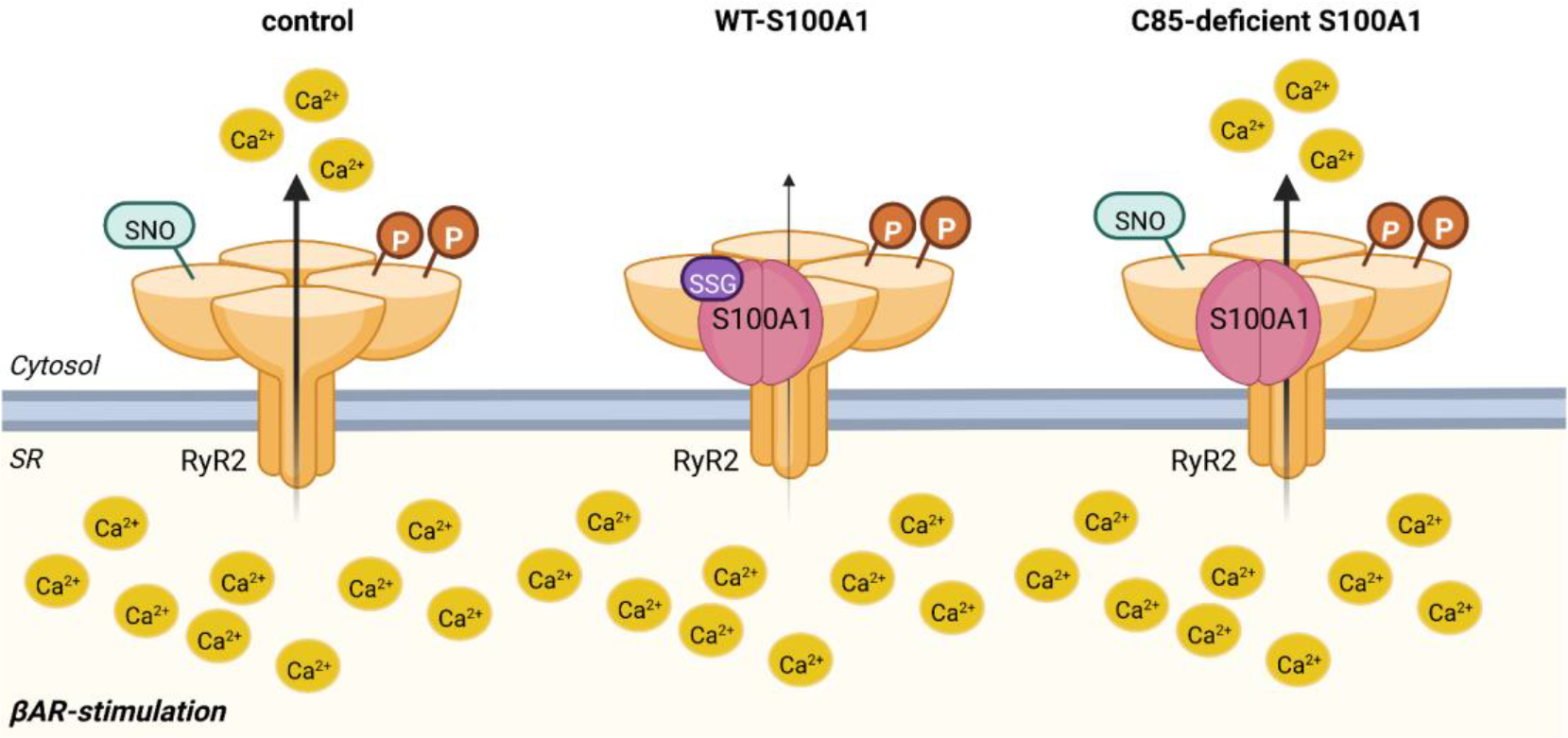
Proposed model for S100A1 regulation of diastolic RyR2 leakage. During βAR stimulation, increased phosphorylation and nitroslyation renders RyR2 leaky, resulting in diastolic Ca^2+^ release. Overexpression of S100A1 prevents RyR2 nitrosylation and subsequently, abrogates diastolic Ca^2+^ leak. Overexpression of S100A1 variants, that lack endogenous glutathionylation sites, fail to attenuate RyR nitroslyation and have no protective effect on diastolic Ca^2+^ leakage. Created with BioRender.com

## 4. Discussion

Taking advantage of engineered S100A1 protein variants, our data show for the first time that the conserved, single C-terminal C85 residue is a key moiety for S100A1’s ability to mitigate diastolic RyR2 dysfunction and, most importantly, to oppose βAR-mediated arrhythmogenic SR Ca^2+^ leakage. Here, the C85 residue appears critical for protecting RyR2 against βAR-induced hypernitrosylation, which is considered to render RyR2s leaky during cardiac relaxation [26, 27, 31]. As such, our study advances our understanding of the molecular mechanism underlying the previously reported antiarrhythmic efficacy of the cardiac EF-hand Ca^2+^ sensor in normal and failing CM *in vitro* and *in vivo* [21, 22].

Our investigation originated from a series of studies that addressed the impact of C85’s ox-PTM on S100A1’s ability to transition from the inactive “apo” to the Ca^2+^ activated “holo” state [10, 12, 14]. In this context, ox-PTM of C85 was demonstrated to significantly improve S100A1’s Ca^2+^ affinity by several orders of magnitude. Using a mBSA, we could confirm that S100A1 is endogenously modified in the adult heart, and specifically, glutathionylated. This is in contrast to a previous report, that suggested that S100A1 is nitrosylated in neuronal cells [12]. However, this observation is in line with the notion that the glutathione concentration is very high in the cell, and thus glutathionylation could be the predominant modification of S100A1 [10].

In CM, ox-PTM of C85 might facilitate a Ca^2+^ activated state of S100A1 already at low diastolic Ca^2+^ levels and subsequently engage with intracellular target proteins [15]. To test this hypothesis and determine if S100A1’s NO-dependent PTM is be required for the molecule to function properly lead us to explore the effect of S100A1 protein variants that lack C85 and ox-PTM in a series of molecular and functional assays making use both of isolated neonatal and adult rat CM.

Analyses of the SR Ca^2+^ leak and diastolic RyR2 function in CM unveiled then the most important result of our study. Imposing significant stress on diastolic RyR2 closure, only WT-S100A1 effectively protected CM against the high number of recurrent Ca^2+^ waves. This is congruent with recent findings demonstrating that WT-S100A1 overexpression protected both normal and failing CM against βAR-mediated exacerbation of proarrhythmogenic diastolic Ca^2+^ waves and after-contractions [21]. In contrast, C85-deficient S100A1 protein variants failed to recapitulate the beneficial diastolic actions of WT-S100A1, particularly upon βAR-stimulation. These results show for the first time that S100A1’s single C85 moiety seems indispensable for the protective effect of WT-S100A1 against diastolic SR Ca^2+^ leak and RyR2 dysfunction.

The hypothesis that S100A1’s single reactive cysteine residue or its NO-dependent PTM might though be required for the molecule to function properly at diastolic Ca^2+^ concentrations prompted us to assess S100A1/RyR2 interaction patterns. Our PLA study performed in resting CM that provide diastolic-like Ca^2+^ concentrations within an intact subcellular microenvironment points toward the notion that C85-deficient S100A1 protein variants can nevertheless interact with the RyR2 to the same extent as WT-S100A1 in the relaxation phase. In agreement with these observations, it has been proposed that Ca^2+^ affinity of S100A1 might also increase upon interaction with the RyR2 and thus, that enhanced Ca^2+^ affinity might be consequence of and not prerequisite for target protein interaction [12, 32]. This is also in line with the idea that S100A1 is an intrinsic disordered protein (IDP) [12]. Although ordered S100A1 regions seem to dominate in the formation of the homodimeric backbone, ox-PTM of C85 is supposed to directly fine-tune the hydrophobic core of S100A1, which restructures the whole protein dimer, exposing sequence fragments ascribed to target binding and subsequent activity modulation [10-12]. This mechanism might unveil a flexible regulatory epitope that could also contribute to diastolic RyR2 activity modulation through a reversible S100A1/RyR2 disulfide bond formation after S100A1 binding to the RyR2 has been stabilized. However, this new hypothesis clearly warrants further validation and detailed structural studies to entail the exact binding mode of S100A1 on the RyR2.

We next focused on the phosphorylation and nitrosylation state of the macromolecular SR Ca^2+^ release channel [24, 25, 27]. The ISO-induced RyR2 phosphorylation at Ser-2808 and Ser-2814 in combination with the more than doubled nitrosylation state might have significantly contributed to diastolic RyR2 dysfunction in the control group [26, 28, 31]. Importantly, inhibition of βAR mediated hypernitrosylation by S100A1 might be the critical link that conveys the protective effect of an enhanced association of the EF-hand Ca^2+^ sensor with the RyR2. These data strongly argue in favor of a critical role for S100A1’s single C85 residue in controlling the RyR2 nitrosylation state, most likely after binding has been established between both molecules.

As RyR2 hypernitrosylation is supposed to render the SR Ca^2+^ channel leaky [31, 33], we propose a model where C85’s ox-PTM of S100A1 specifically modulates diastolic RyR2 activity by controlling the channel’s redox-state. This is further supported by the finding that systolic performance by overexpressed C85-deficient S100A1 protein variants was indistinguishable from WT-S100A1 protein.

Hence, this prompts the question how the single cysteine might contribute to S100A1-mediated RyR2 regulation? Aiming to corroborate a direct link between NO-regulation and S100A1, we focused our effort on NOS1, which is know the regulate RyR2 activity and SR Ca2+ handling [28, 29]. However, this link could not be established, which argues against a role for S100A1 in the control of overall CM redox homeostasis. Alternatively, spatiotemporal changes in NO and ROS have been proposed to regulate nitrosylation and subsequently activity of the RyR2 [28, 34]. However, in order to better understand this concept, improvement of experimental methods to detect fluctuating NO levels are clearly needed.

In summary, dissecting S100A1’s molecular structure-function relationship in CM, our data indicate for the first time that the c-terminal conserved single cysteine C85 is inevitable for WT-S100A1’s antiarrhythmic but dispensable for its inotropic efficacy in CM. Our data strongly support the notion that the protective effect of S100A1 against diastolic RyR2 dysfunction might rely on the control of the RyR2 nitrosylation state and more specifically unveil a critical role of the C85 moiety in abrogating RyR2 hypernitrosylation. Therefore, we propose a simplified 2-step model where C85 seems dispensable for S100A1/RyR2 interaction but indispensable for beneficially modulating diastolic RyR2 activity. This effect could be due to the IDP properties of WT-S100A1 that requires the presence and ox-PTM of C85.

Our novel results doubtlessly advance our understanding of the molecular mechanisms underlying the beneficial pleiotropic actions of WT-S100A1 in the heart. Continued research is now needed to delineate S100A1 binding epitopes with the RyR2 and its potential involvement in the spatially confined SR nitrosylation/denitrosylation cycle. This might aid a deepened understanding of the molecular S100A1/RyR2 liaison given the therapeutic potential of S100A1 in heart failure.

## Statements & Declarations

### Author Contributions

AS, JK, AS, SS, AJ and JR performed the experiments; AS, JR and PM designed the experiments and AS, MB and JR analyzed the data; AS, JR, PM and wrote the manuscript; HG provided research support. All authors read and approved the final manuscript.

### Data availability

Additional data supporting findings of this study can be found in the supplementary material.

